# Energetic and structural features of SARS-CoV-2 N-protein co-assemblies with nucleic acids

**DOI:** 10.1101/2021.02.08.430344

**Authors:** Huaying Zhao, Di Wu, Ai Nguyen, Yan Li, Regina C. Adão, Eugene Valkov, George H. Patterson, Grzegorz Piszczek, Peter Schuck

## Abstract

Nucleocapsid (N) protein of the SARS-CoV-2 virus packages the viral genome into well-defined ribonucleoprotein particles, but the molecular pathway is still unclear. N-protein is dimeric and consists of two folded domains with nucleic acid (NA) binding sites, surrounded by intrinsically disordered regions that promote liquid-liquid phase separation. Here we use biophysical tools to study N-protein interactions with oligonucleotides of different length, examining the size, composition, secondary structure, and energetics of the resulting states. We observe formation of supramolecular clusters or nuclei preceding growth into phase-separated droplets. Short hexanucleotide NA forms compact 2:2 N-protein/NA complexes with reduced disorder. Longer oligonucleotides expose additional N-protein interactions and multi-valent protein-NA interactions, which generate higher-order mixed oligomers and simultaneously promote growth of droplets. Phase separation is accompanied by a significant increase in protein secondary structure, different from that caused by initial NA binding, which may contribute to the assembly of ribonucleoprotein particles within molecular condensates.

## Introduction

The ongoing COVID19 pandemic is caused by the severe acute respiratory syndrome coronavirus 2 (SARS-CoV-2), and concerted research into the molecular mechanisms of this virus and its interactions with the host cells has been undertaken worldwide with the aim to develop vaccines and therapeutics. In particular, unprecedentedly rapid progress has been made in the analysis of the viral spike protein which is critical for inhibition of viral entry and for the development of immunity. Extensive work in this area by many laboratories has led to vaccines based on spike protein antigens, and to antibody therapeutics (Hansen et al., 2020; Krammer, 2020). Viral replication is a second point of attack (Chen et al., 2020b; Hillen et al., 2020; Wang et al., 2020), most prominently with the use of nucleoside analogues interfering with viral polymerase function (Pruijssers et al., 2020). Recently emerging spike mutant strains (Rambaut et al., 2020) stress the importance of parallel and novel therapeutic approaches. For other viral diseases, various aspects of viral assembly have previously emerged as additional antiviral targets (Baines, 2011; Hurt et al., 2011; Keil et al., 2020; Spearman, 2016). Unfortunately, our knowledge about SARS-CoV-2 assembly is still very limited, and efforts in many laboratories are underway to elucidate this key step in the viral life cycle. Extending earlier structural studies of coronaviruses (Gui et al., 2017; Macnaughton et al., 1978), it has recently emerged that SARS-CoV-2 virions contain on average 38 ribonucleoprotein particles (RNPs), each consisting of a ≈14-15 nm diameter cylindrical arrangement of pillars composed of viral nucleocapsid (N) protein, which are thought to package the large viral genome of ssRNA in a nucleosome-like fashion (Klein et al., 2020; Yao et al., 2020). Here we focus on the initial stages of the RNP assembly by the viral N-protein.

SARS-CoV-2 belongs to the family of betacoronaviruses and its N-protein is highly conserved (Tilocca et al., 2020; Zhou et al., 2020). In particular, it is 89.7 % homologous to that of SARS-CoV-1 (Kang et al., 2020), which has been studied since the 2002/2003 outbreak, and much of what has been learned about SARS-CoV-1 N-protein is thought to apply similarly to SARS-CoV-2 (Chang et al., 2014; Masters, 2006, 2019). N-protein from coronavirus is a basic multi-domain structural protein consisting of two folded domains at the N-terminus (NTD) and C-terminus (CTD) flanked and linked by intrinsically disordered amino acid stretches (Chang et al., 2006) (**Figure 1A**). It has a molecular weight of 45.6 kDa but forms a tightly bound dimer in solution mediated by interactions in the CTD. N-protein is the most abundant viral protein and presents as a major antigen, although a strong immune response against N-protein is associated with comparatively poor protection (Atyeo et al., 2020; Yasui et al., 2008). It is multi-functional (McBride et al., 2014), and intracellular binding partners include several proteins involved in RNA processing, and in stress granule regulation (Gordon et al., 2020; Nabeel-Shah et al., 2020), but its most obvious role is structural, in binding viral ssRNA and assembling RNPs. This is accomplished with the help of nucleic acid (NA) binding sites at both NTD and CTD of the highly basic N-protein, as well as in its disordered regions, which bind NAs in a largely sequence independent fashion (Chang et al., 2009; Takeda et al., 2008). Additionally, N protein interacts with the endodomain of viral M-protein, which is thought to anchor RNPs to the viral membrane and may play a role in the recognition of the viral packaging signal (He et al., 2004a; Masters, 2019).

**Figure 1.**
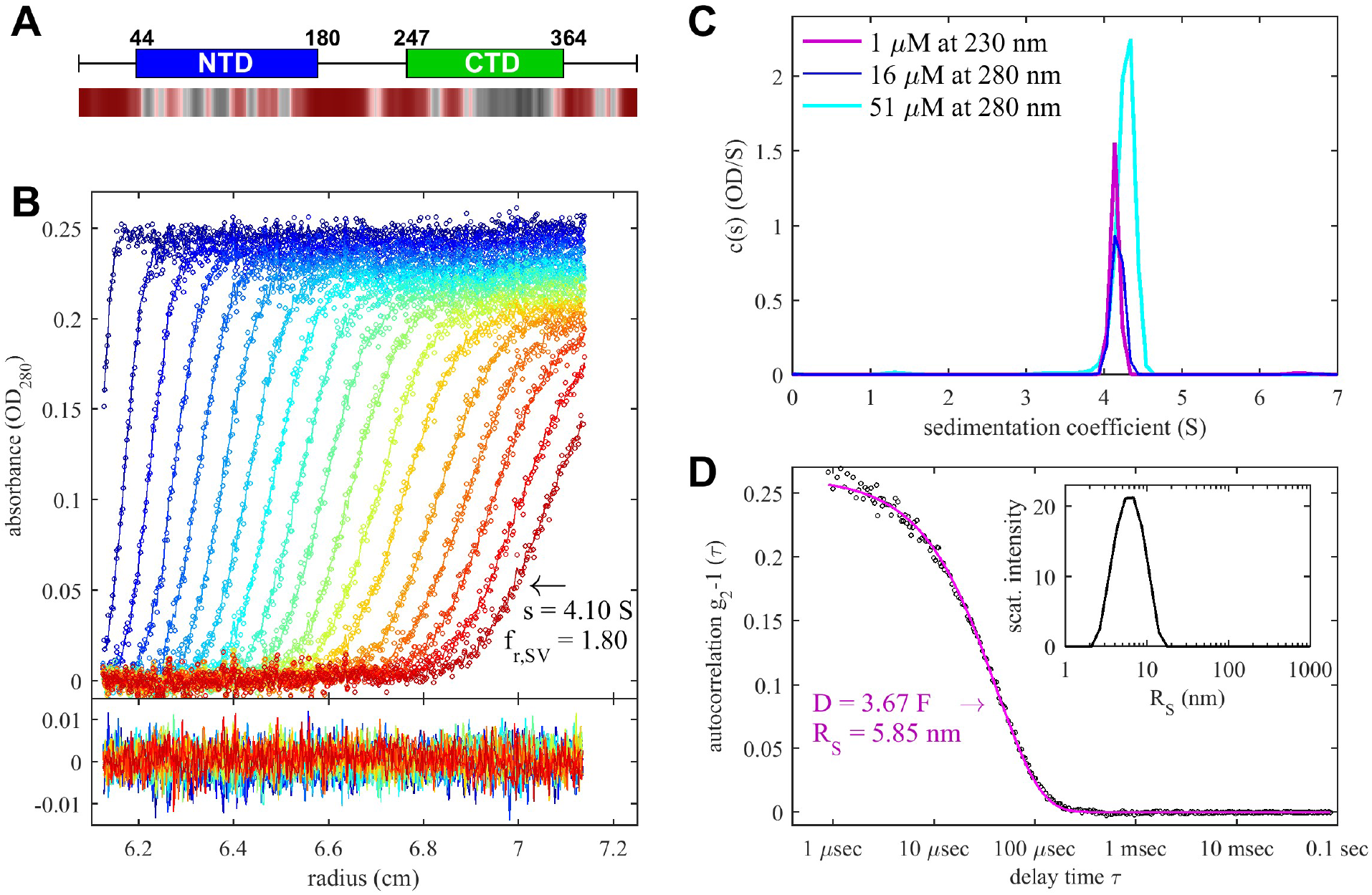
SARS-CoV-2 N-Protein Forms a Stable Dimer in Solution. (A) Top: Organization of N-protein primary structure with N-terminal domain (NTD) and C-terminal domain (CDT) flanked and linked by disordered stretches (Dinesh et al., 2020; Zinzula et al., 2020). Bottom: Propensity of residues to promote LLPS calculated by FuzzDrop, showing in red droplet-promoting regions with p_DP_ >0.6 (Hardenberg et al., 2020). (B) Top: Boundaries of 5 μM N-protein sedimenting at 50,000 rpm in SV-AUC (symbols) and *c*(*s*) sedimentation coefficient distribution model (lines; color temperature indicating scan time). Bottom: Residuals of the fit. (C) Sedimentation coefficient distributions at different concentrations of N-protein. (D) DLS autocorrelation data of 3 μM N-protein (symbols) with single discrete species fit (line). Result from a Stokes radius distribution model is shown in the inset.

Structures and dynamics of both CTD and NTD, isolated or in complex with NA, have been studied and reveal significant molecular details (Ahamad et al., 2020; Caruso et al., 2020; Chen et al., 2007; Dinesh et al., 2020; Kang et al., 2020; Takeda et al., 2008; Zinzula et al., 2020). Intriguingly, it was reported recently by several laboratories that N-protein undergoes liquid-liquid phase separation (LLPS) (Carlson et al., 2020; Cascarina and Ross, 2020; Chen et al., 2020a; Cubuk et al., 2020; Iserman et al., 2020; Jack et al., 2020; Lu et al., 2020; Perdikari et al., 2020; Savastano et al., 2020; Wang et al., 2021; Zhao et al., 2020). Such demixing is commonly mediated by ultra-weak but highly multi-valent interactions (Alberti et al., 2019; Vernon et al., 2018). LLPS was proposed to aid in efficient replication through condensates that recruit the RNA polymerase, and/or to interfere with stress granules function, though it is also likely that a dense phase containing N protein and viral RNA can promote assembly due to the high local concentrations. Alternate functions may be promoted by different degree of N-protein phosphorylation (Fung and Liu, 2018; Wu et al., 2009).

On a molecular level, much of the assembly pathway remains elusive: we do not know how the assembly is initiated, what are the molecular building blocks of RNP, and what are the driving forces stabilizing higher-order complexes in dilute phases or in macromolecular condensates. Likewise, it is unknown what are the intermediate assembly states, their stoichiometry, conformation, and energetics, on the path leading up to complete RNPs. Clearly, they must result from as of yet unknown – and potentially druggable – protein-protein and protein-RNA interfaces.

To address these questions, we apply a suite of hydrodynamic, spectroscopic, and calorimetric biophysical methods to study size, composition, solution structure, and thermodynamic stabilities of full-length SARS-CoV-2 N-protein by itself and in complexes with oligonucleotides of different lengths. This reveals architectural and energetic aspects of the initial steps of RNP assembly, and offers new insights in the dynamics of higher-order oligomers, clusters, and biomolecular condensates.

## Results

### Solution conformation of SARS-CoV-2 N-protein and energetics of dimerization

The characterization of N-protein self-association is a foundation for ascertaining NA ligand-linked cooperativity in assembly. We first applied sedimentation velocity analytical ultracentrifugation (SV-AUC) to measure sedimentation coefficient distributions *c*(*s*) that report on solution states of N-protein and its complexes. **Figure 1B** shows a representative set of raw sedimentation boundaries for 5 μM N-protein at 20°C under our standard experimental condition of 12 mM KH_2_PO_4_/Na_2_HPO_4_, 3 mM KCl, 10 mM NaCl, pH 7.4. Via the Svedberg relationship (Svedberg and Pedersen, 1940), sedimentation and boundary diffusion implies a mass estimate from SV-AUC of 87.3 kDa. **Figure 1C** shows the concentration-dependence of the sedimentation coefficient distributions *c*(*s*), highlighting the monodispersity of the protein. Complementary experiments were carried out by dynamic light scattering (DLS), which yields autocorrelation functions that can be described well by a single class of particles with Stokes radius of 5.85 nm (**Figure 1D**). When the sedimentation coefficient of 4.1 S from SV-AUC is combined with the diffusion coefficient from DLS a molecular weight of 96.2 kDa is obtained, which is within error of the 94 kDa of a dimer. This shows the N-protein dimerizes with a highly non-compact solution structure (f/f_0_ = 1.82) as expected from SARS-CoV-1 (Yu et al., 2005), and consistent with the SAXS scattering data of SARS-CoV-2 by Zeng et al (Zeng et al., 2020).

Notably, we observe that at all concentrations (1 – 51 μM) the SV-AUC signals observed by both absorbance and refractometric detection were at the expected magnitude. This shows the absence of LLPS under these conditions, since LLPS would create dense phases that rapidly sediment and be discernable indirectly from loss of remaining sedimentation signal amplitudes at the higher concentrations (see below). Also, both SV-AUC and DLS revealed little indications of higher-order oligomers, traces of which both techniques would detect with high sensitivity. The slight increase in sedimentation coefficient at the highest concentration is consistent with transient dimerization arising from ultra-weak protein-protein interactions with a best-fit K_D_ of 0.76 mM, reminiscent of SARS-CoV-1 N-protein (Yu et al., 2005). Likewise, very little dissociation of the dimer is observed within the experimentally accessible concentration range (**Figure 1BC**), suggesting K_D_ < 10 nM. No change in association state was observed across a large range of temperature, assessed by SV-AUC a 10°C (data not shown), DLS up to ≈45°C (below). We also examined the ionic strength dependence of dimerization, but no significant difference in oligomeric state was observed in SV-AUC with 10 mM NaCl, 150 mM NaCl, 300 mM, or 600 mM NaCl.

To quantify the dimer affinity we aimed to use nanomolar concentrations, however, this was hampered by the stickiness of the N-protein to the AUC sample cell. We found this can be overcome by using PBS buffer with 0.005% of the surfactant P20. Under this condition, SV-AUC with far-UV detection showed substantial dissociation below 50 nM, with a population of a ≈2 S monomer species (**Figure S1A**) and an estimated K_D_ of 29 nM (95%CI [5.2, 340] nM). Similarly, after fluorescently labeling N-protein with Dylight488 *via* amine coupling, we observe reversible dissociation with a K_D_ of 1.2 nM (< 4.5 nM within 95%CI) (**Figure S1C**).

To assess N protein secondary structure we carried out circular dichroism spectroscopy (CD) in different conditions (**Figure S2** and **Figure 3C** below). Overall, the spectra are consistent with those measured recently by Zeng et al (Zeng et al., 2020). Even though they do not exhibit the canonical features of well-folded proteins, the very strong negative ellipticity at 200 nm shows significant disordered fractions. Interestingly, addition of surfactant or the fluorescent tag shows slight alterations in the protein (**Figure S2**). This raises the possibility that the intrinsically disordered regions may be in a delicate conformational balance susceptible to small structural perturbations or changes in solvation. Therefore, we omitted the use of fluorescence tags and surfactant in the following.

In summary, at neutral pH, moderate ionic strength and temperature, SARS-CoV-2 N-protein is highly extended in solution, significantly disordered, and in a tight monomer-dimer equilibrium. There is no evidence of higher-order oligomers, except for rapid ultra-weak interactions, and no LLPS across a wide range of conditions explored here.

### Length-dependent mode of short oligonucleotides binding to SARS-CoV-2 N-protein dimer

In view of the fact that N-protein forms large ribonucleoprotein particles, it seems plausible that N-protein self-association is coupled to NA binding in a co-assembly process. SARS-CoV-1 N-protein has multiple binding sites that can interact with single-stranded RNA or DNA in a broadly sequence independent fashion (Chang et al., 2014; Chen et al., 2007). We aimed to dissect N-protein/NA binding and the onset of co-assembly using short oligonucleotide probes of different lengths, including GT, (GT)_2_, T_6_, T_10_, and T_20_.

In preliminary experiments we observed the expected strong ionic strength dependence of NA binding. We selected standard buffer condition with 10 mM NaCl and pH 7.4 to populate complex states on a concentration scale allowing for biophysical characterization, with selected control experiments in phosphate buffered saline (PBS) for comparison. SV-AUC is uniquely capable of resolving different free and complex species in solution by size, while simultaneously reporting on rapidly sedimenting dense fractions after LLPS. The Rayleigh interference optical system reports mostly on protein, while absorbance signals at 280 nm, 260 nm, and 230 nm report on NAs and proteins in distinctive ratios, revealing species’ stoichiometries. To this end, we have determined molar signal increments separately for N-protein and for all oligonucleotides. Thus, binding of oligonucleotides can be ascertained in SV-AUC in multiple ways: 1) from depletion of the population of slowly sedimenting unbound NA; 2) from shifts in the sedimentation coefficient of the protein (or complex) due to added mass and/or altered protein oligomeric state; 3) from increased ratio of absorbance at 260 nm vs 280 nm associated with the sedimentation boundary of the protein/NA complex(es); and 4) for NA partitioning into the rapidly sedimenting dense phase after LLPS, thereby creating loss of the total protein and NA signal remaining in solution after obligatory initial rapid depletion of all dense phase droplets at the start of centrifugation.

For the smallest oligonucleotide, GT, there was no indication of binding by any of these measures (**Figure 2A**). By contrast, for (GT)_2_ a small concentration-dependent shift in the protein dimer s-value from 3.9 S to 4.1 S was observed with increasing NA ligand, accompanied by a significantly enhanced 260 nm signal co-sedimenting with the main boundary (**Figure 2B**). (Binding of (GT)_2_ to N-protein was confirmed independently by differential scanning fluorometry; see below.) With 5-fold molar excess of (GT)_2_ we determined a molar ratio of only 1:1 N-protein/(GT)_2_, which shows that the average complex in solution is an N-protein dimer with two (GT)_2_ oligonucleotides. However, we were unable to carry out experiments at high enough concentrations to approach saturation even at 10-50 μM, indicating K_D_ above 10 μM in our low salt buffer. By contrast, binding was completely abrogated at 137 mM NaCl in PBS (**Figure S3A**).

**Figure 2.**
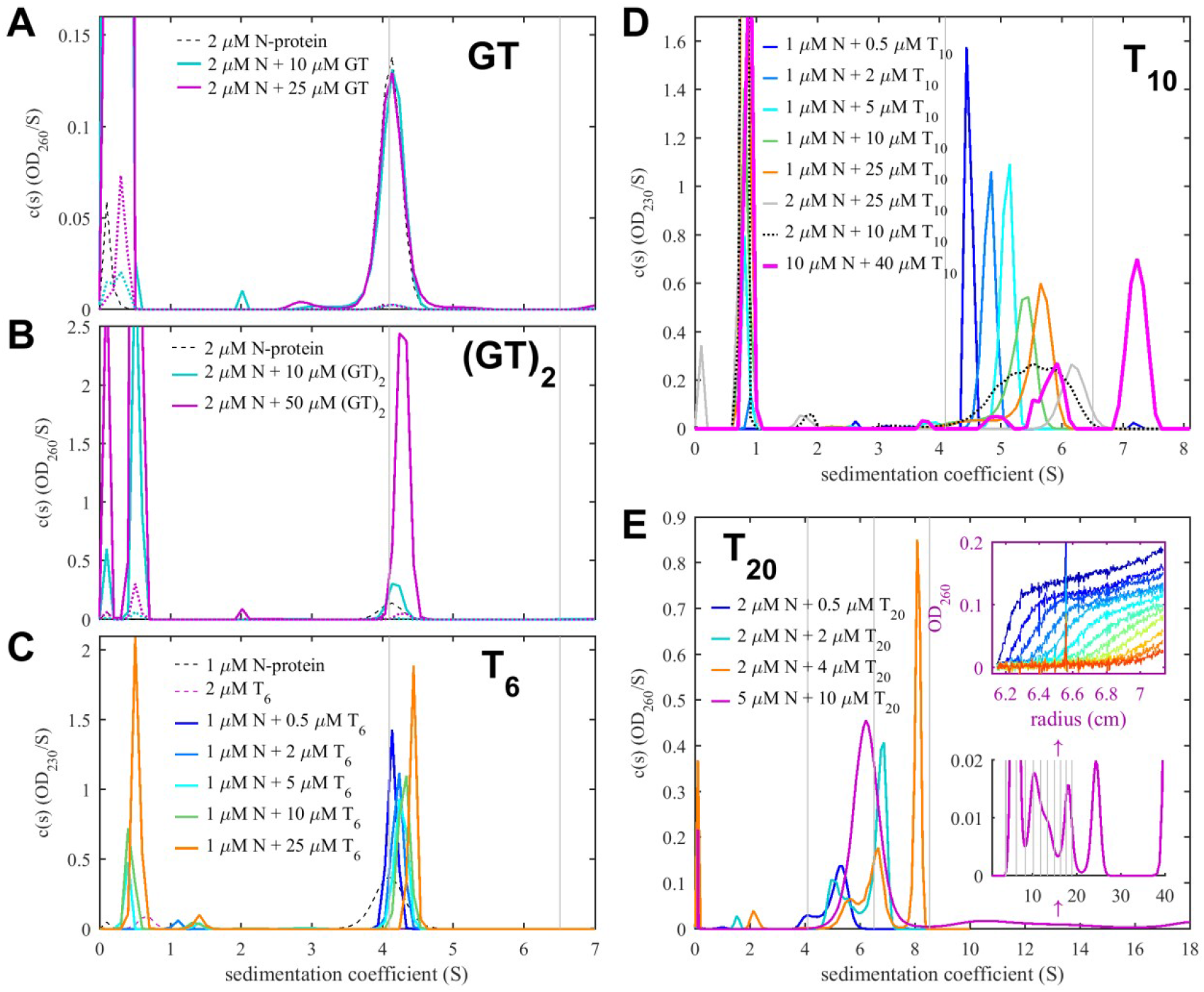
NA-Linked Self-Association of N-protein Revealed in SV-AUC. Shown are sedimentation coefficient distributions c(s) obtained from mixtures indicated. As a visual guide, thin vertical grey lines indicate s-values of unliganded N-protein dimer and its oligomers based on the M^2/3^ hydrodynamic scaling law. (A) No significant binding is observed for GT, as indicated by lack of change in sedimentation coefficient distribution. Dotted lines are 50-fold magnified traces. (B) (GT)_2_ binding can be observed from increase in the 260 nm signal co-sedimenting with N-protein (C) Enhanced 260 nm signal co-sedimenting with N-protein, decreasing free T_6_ signals, and increased s-values of the protein sedimentation coefficient show significant binding, with weighted-average s-values in **Figure 3A**. (D) Continuous increase in sedimentation coefficient with T_10_ concentration shows N-protein dimer-dimer interactions forming rapidly exchanging oligomers. (E) Efficient binding of T_20_ leads to oligomers larger than tetramers and, at high concentrations, skewed leading plateaus in the sedimentation boundaries (upper inset) corresponding to a ladder of oligomers extending to > 40 S (lower inset). LLPS is revealed by cloudiness of the T_20_/N-protein mixtures prior to sedimentation and loss of signal in the remaining sedimentation boundaries.

The slightly larger 6nt oligonucleotide T_6_ exhibited clear binding to N-protein at low μM concentrations (**Figure 2C**) with complex *s*-values increasing up to 4.91 S. At high protein concentrations with large excess of T_6_, absorbance signal amplitudes indicate a stoichiometry of two T_6_ per N-protein dimer. It is noteworthy that there was no indication of either dissociation of N-protein dimers or formation of higher-order oligomers up to concentrations of 10 μM N-protein in the presence of either GT, (GT)_2_, and T_6_, suggesting the protein dimers to be the building blocks for all observed N-protein assemblies.

The robust, saturable binding of T_6_ permits more quantitative analysis of binding affinity. To this end, we carried out titration series in SV-AUC at different concentrations of N-protein from 1 to 15 μM, and extracted isotherms of weight-average *s*-values (**Figure 3A**). In addition, we carried out isothermal titration microcalorimetry (ITC) experiments titrating T_6_ into N-protein (**Figure 3B**), followed by SV-AUC control experiments of the final cell contents to verify the absence of LLPS and aggregation, and hydrodynamically confirming complex formation. SV-AUC and ITC titration isotherms were jointly fit in global multi-method modeling, yielding an excellent fit with two equivalent, non-cooperative binding sites per N-protein dimer, with microscopic *K_D_* = 0.6 (95% CI 0.1 – 0.7) μM, arising from both favorable enthalpy ΔH = −6.4 (95% CI −7.7 to −5.1) kcal/mol and favorable entropy of TΔS = 2.4 (95% CI 0.2 – 5.3) kcal/mol. We also attempted a more detailed SV-AUC analysis including titration series at lower N-protein concentrations of 0.5 μM in the presence of 0.005% P20, which resulted in a best-fit model with slight negative cooperativity of sites with macroscopic *K_D_*-values of 0.36 μM and 1.8 μM (not shown).

**Figure 3.**
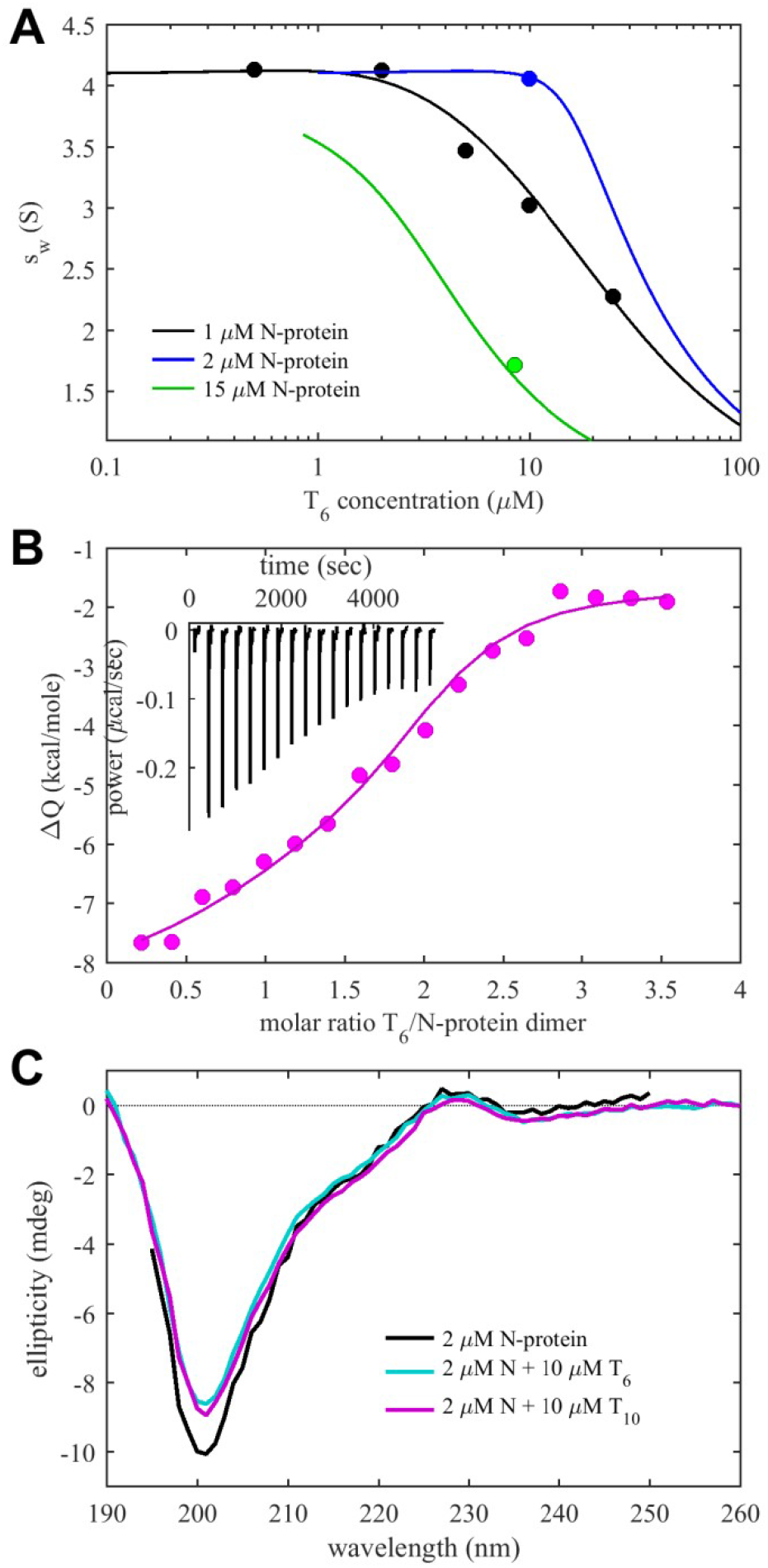
Conformational Change and Energetics of Oligonucleotide Binding. (A) Weight-average sedimentation coefficients s_w_ from SV-AUC titration series of N-protein with T6 (symbols) and best joint fit with ITC data (lines). (B) ITC titration isotherm titrating 200 μM T_6_ into 24 μM N-protein (symbols) and best joint fit with s_w_ isotherms in Panel A (line). The inset shows the baseline corrected power trace. (C) Circular dichroism spectra of N-protein and its complexes with oligonucleotides T_6_ and T_10_.

CD spectra acquired in the presence of saturating T_6_ are shown in **Figure 3C**. As mentioned above, the spectra exhibit a strong negative ellipticity at 200 nm associated with disordered regions. Interestingly, under conditions where the N protein dimer is liganded by two T_6_ oligonucleotides, the negative signal is significantly reduced. This suggests a reduction of disorder of the polypeptide chain due to NA binding. The spectra would be consistent, for example, with formation relaxed anti-parallel β-sheets (Micsonai et al., 2015).

### Higher oligomerization of SARS-CoV-2 N-protein dimer in complex with longer oligonucleotides

New behavior was observed in binding experiments with the longer oligonucleotides T_10_ and T_20_. For the decanucleotide T_10_, the sedimentation coefficient distributions measured at various molar ratios are shown in **Figure 2D**. Notably, already at the lowest concentration of 1 μM protein with 0.5 μM T_10_, based on the characteristic 260 nm absorbance traces, there is virtually no T_10_ left unbound. This demonstrates stronger binding for T_10_ than T_6_, with K_D_ <0.1 μM. With higher decanucleotide concentrations, complex peaks in the sedimentation coefficient distribution show a shift to higher s-values, while the boundaries exhibit excess broadening with time. Such behavior is characteristic for reaction boundaries where all free and complex species are in a rapid exchange and migrate as a coupled system with a sedimentation coefficient reflecting the time-average of mixed assembly states (Schuck, 2010; Schuck and Zhao, 2017).

Particularly interesting is the experiment at the highest concentrations with 10 μM N-protein and 4-fold excess of T_10_ (bold magenta line in **Figure 2D**). Here, we observed the highest *s*-value of ≈7.5 S, almost twice that of the N-protein dimer. Theoretically, a dimer ligated by 4 copies of T_10_ could assume an *s*-value of 7.5 S only if it had a frictional ratio of < 1.15, which would correspond roughly to a hydrated compact sphere – but this geometry that can be excluded based on N-protein organization. Therefore, we can deduce that T_10_ promotes dimer-dimer interactions forming tetramers and possibly higher oligomers. In this experiment, the signal ratios of the observed complex reveal an average molar ratio of 1:2 (or 4 NA per dimeric N-protein unit), which is twice that of T_6_. Furthermore, in contrast to experiments with T_10_ added to lower protein concentrations, cloudiness in the solution developed within seconds of mixing this sample, suggesting the formation of large clusters or droplets from LLPS, as is examined in more detail below. We found these to be largely reversible upon dilution (data not shown). As mentioned above, dense phase droplets will sediment instantly and the measured SV-AUC data display the properties of the dilute phase at the phase boundary of LLPS.

Complexes with similar N/T_10_ molar ratios were measured for lower total protein concentrations (2 μM) in larger excess of NA, with a value of 1:2 in 5-fold excess T_10_, and even 1:2.5 in 25-fold excess T_10_, respectively. This molar ratio implies binding of at least two NA binding sites are present on each N-protein protomer and are engaged with different T_10_ molecules, and possibly additional binding at a weaker third site. CD experiments under these conditions show reduced negative ellipticity at ≈ 200 nm, virtually identical to those observed in the presence with T_6_ ligand, again indicating gain of structure in the polypeptide chain (**Figure 3C**).

The fact that the *s*-values of N/T_10_ complexes strongly increase with protein concentration reveals the formation of larger assemblies engaging multiple N-protein dimers. Taken together with the observation of an approximately constant molar ratio of 1:2 (for all mixtures with excess T_10_ up to 25-fold), a picture arises where 4-fold liganded dimers present units that self-associate in an N-protein concentration-dependent manner with and effective *K_D_* for dimer-dimer association in the low micromolar range.

The longest oligonucleotide we studied is the eicosamer T_20_. SV-AUC data for the mixtures of N-protein with the T_20_ are shown in **Figure 2E**. First, at low concentrations of 2 μM N-protein with only 0.5 μM T_20_ (blue) all oligonucleotide was co-sedimenting with the protein, consistent with the high affinity binding seen already with T_10_. Also, there is little free protein despite its 4-fold molar excess, with the majority of protein sedimenting in a complex at ≈5.2 S, distinctly faster than would be expected solely by added mass of T_20_ (6.0 kDa). This shows that on average more than one N-protein dimer is engaged in a complex with a single T_20_. Noting the continuously increasing sedimentation coefficient with increasing N-protein concentration, we can conclude this interaction forms a reaction boundary of a dynamic system, which occurs with complex life-times less than ≈1,000 sec. High s-values and creation of higher N-protein assemblies were already observed in complexes with T_10_, but were associated with high saturation of NA sites on the N-protein dimer; by contrast, a single T_20_ can already crosslink or scaffold N-protein dimer, as schematically illustrated in **Figure 4**.

**Figure 4.**
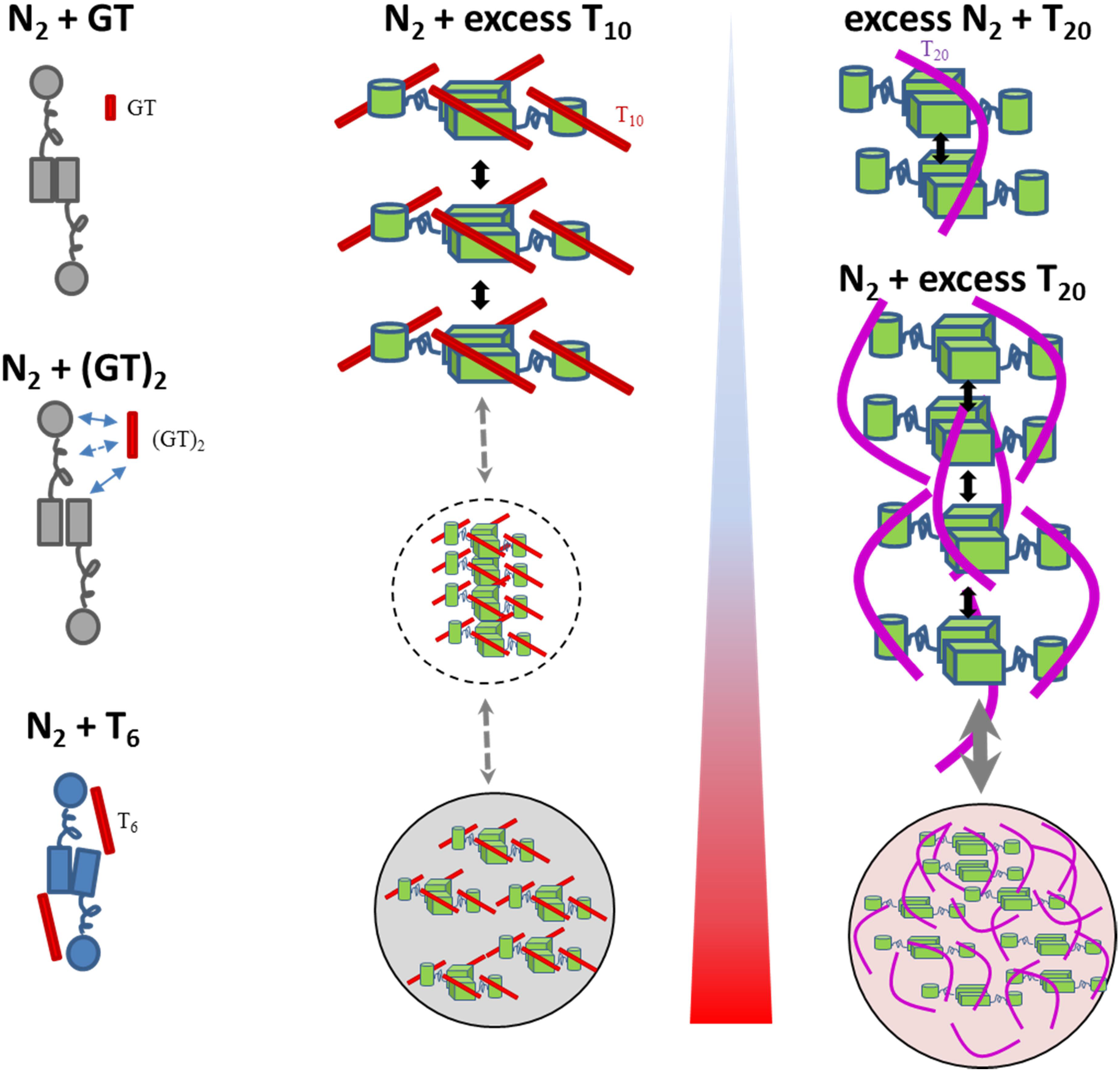
Schematics of co-assembly with different oligonucleotides. The cartoon depiction highlights oligomeric architecture and interactions, without detail on protein structures. Domain organization is sketched after published cryo-EM and SAXS data (Chang et al., 2009; Zeng et al., 2020; Zinzula et al., 2020), schematically highlighting disordered linkers and different N-protein conformations in different colors, as indicated by changes in CD spectra. Arrows depict interactions and assembly pathways including clusters and droplets encircled by dotted and dashed lines. The blue to red shaded triangle depicts temperature driven processes.

In all mixtures with 2 μM or higher N-protein in the presence of equimolar or higher T_20_ concentrations, complex species with velocities corresponding to tetramers and hexamers, are observed, and formation of rapidly sedimenting dense phases from LLPS can be deduced from cloudiness after mixing and depletion of macromolecules in the range of measurable sedimentation coefficients (< ≈100 S). The species remaining in the measured dilute phase at the LLPS phase boundary depend on solution composition. They exhibit the characteristic multimodal sedimentation pattern of reaction boundaries arising from oligomeric states in rapid exchange (**Figure 2E**). For example, a mixture of 2 μM protein with equimolar T_20_ (green) shows a typical bimodal boundary, and from decomposition of absorbance and interference signals, we measure a molar ratio of 3.4:1 protein:T_20_ in the slower (≈5S) peak, and a much lower molar ratio of 1 – 1.5:1 in the faster (≈7S) peak. This shows that more than one T_20_ per N-protein dimer can dynamically cross-link dimers into larger mixed oligomers. Similarly, a loading mixture of 2 μM N-protein and 4 μM T_20_ (orange) exhibits sedimentation boundaries with ≈6.5 S and ≈8 S, and spectral decomposition of the latter leads to a molar ratio of ≈1:1. 8 S exceeds the *s*-value expected of an N-protein tetramer, which suggests that higher oligomerization is promoted by T_20_. Finally, in experiments with loading concentrations of 5 μM N-protein with 10 μM T_20_ we can discern a ladder of even larger species in solution (**Figure 2E inset**), including complexes in excess of 20 S. Such a pattern would be expected of an indefinite self-assembly reaction leading up to LLPS. In high salt conditions (PBS) binding of T_20_ is substantially weaker, and lower s-values are observed suggesting the absence of species higher than N-protein tetramers (**Figure S3B**), highlighting the electrostatic contributions to the higher-order oligomerization. The schematics **Figure 4** summarizes different complex architectures and interaction modes of all oligonucleotides.

### SARS-CoV-2 N-protein can form clusters and droplets in mixtures with oligonucleotides

It has been previously reported that N-protein can phase separate with RNA (Chen et al., 2020a; Cubuk et al., 2020; Savastano et al., 2020), and N-protein scores very high for predicted LLPS supported by residues outside its folded domains (**Figure 1A**). LLPS is generally driven by weak but highly multi-valent macromolecular interactions (Alberti et al., 2019). These are inseparably coupled to the co-assembly mechanisms and interactions observed in dilute solution and their reaction products that may facilitate or oppose LLPS.

As described above, at room temperature we have seen cloudiness develop upon mixing μM N-protein with oligonucleotides, increasing in propensity for LLPS with increasing strength and number of protein-protein and protein-NA interactions: Under our conditions, cloudiness was observed neither for N-protein alone, nor in mixtures with T_6_, where there is saturable nucleic acid binding but no significant higher-order oligomerization. However, binding of T_10_ supports weak dimer-dimer interactions, and we observed LLPS at the highest concentration of N-protein with large excess of T_10_. In SV-AUC, the depletion of the sum of free and oligomer-bound NA and protein allows us to determine the average composition of the rapidly sedimenting material. For example, the mixture of 10 μM N-protein and 40μM T_10_ creates a dense phase comprising 2.5 μM of the N-protein and 11 μM of T_10_, thus accommodating 4-fold excess of T_10_ over protein. Finally, T_20_ can additionally cross-link multiple N-protein dimers at low concentrations, and we see cloudiness already at much lower concentrations of ≥ 2 μM N-protein with equimolar or more T_20_. At the low concentrations used, only a small fraction of material was observed to enter the dense phase, which limits the precision for determining its protein/T_20_ molar ratio, with an estimated range of 1:2 – 1:5.

To examine the size of the rapidly sedimenting particles, we turn to DLS and widefield light microscopy. The effects of T_6_ and T_10_ on the size distribution of N-protein are shown in **Figure 5AB** (cyan and orange). In addition to the small peak at the Stokes radius of ≈6 nm of the free N-protein dimer, the scattering signal is dominated by particles at ≈100 nm, with those of T_6_ noticeably fewer and smaller than those of T_10_. Small additional fluctuations on the msec timescale would correspond to 1 – 10 μm particles but exceed the size range of hydrodynamic interpretation of DLS. Since the scattering intensity grows strongly with particle size, the number of the large particles is actually very small. (A back-of-the-envelope estimate suggests particles at 10fold radius produce the same signal at 1/1000th of mass.) Widefield microscopy did not show significant droplets for N-protein alone under these conditions (data not shown). By contrast, formation of droplets outside the range for DLS, on the order of ≈1-2 μm, can be observed in light microscopy for mixtures of 10 μM N-protein with 40 μM T_10_, and in 5 μM N-protein in the presence of 10 μM T_20_ (**Figure 5C**). These droplets were observed to readily dissolve upon dilution (**Figure S4**).

**Figure 5.**
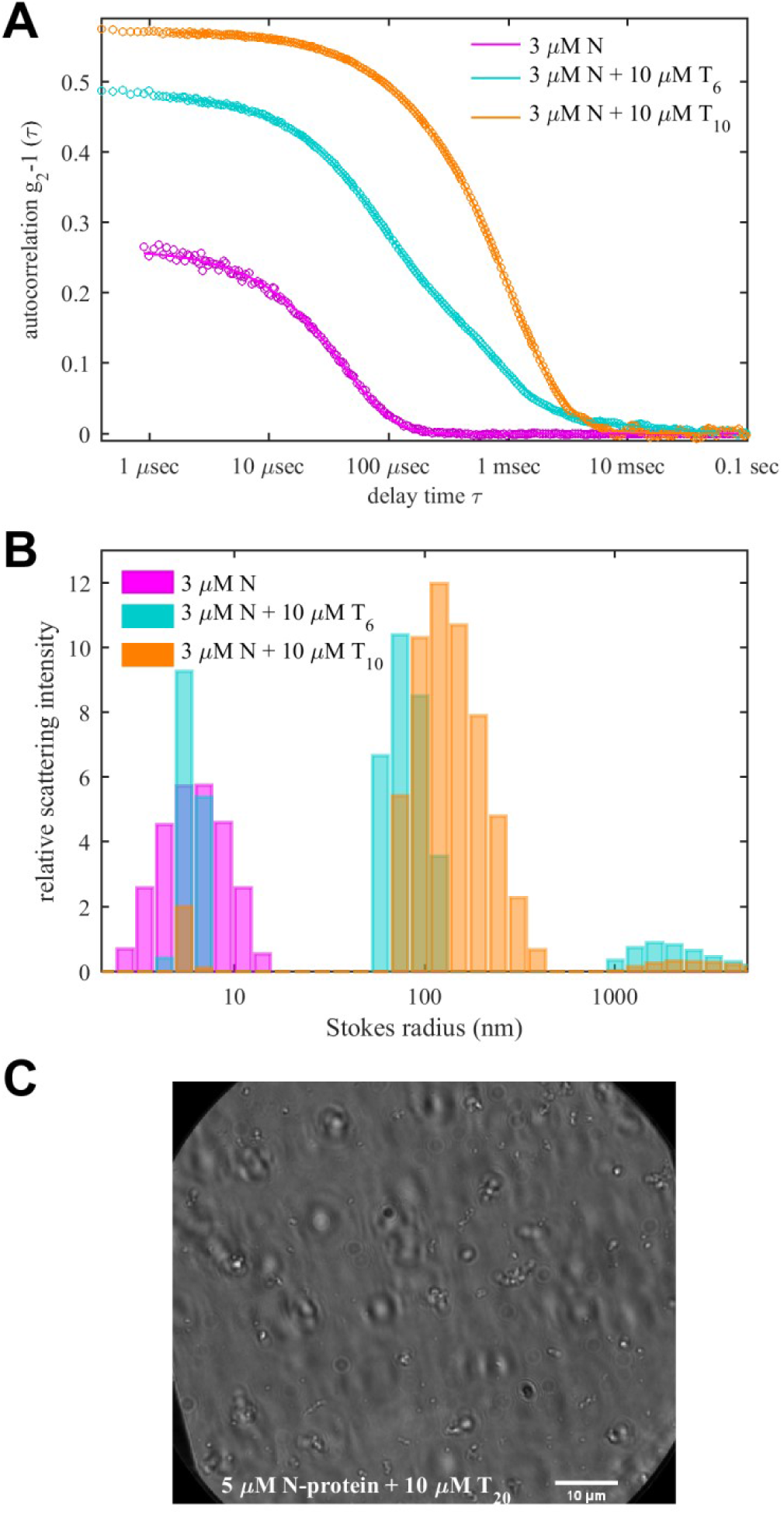
Particle Sizes of Clusters and Droplets. (A) DLS autocorrelation functions of N-protein in the presence of T_6_ and T_10_ oligonucleotides. (B) Stokes radius distributions in scattering intensity units calculated from the data in (A). (C) Widefield microscopy image of N-protein with T_20_.

### Energetic and structural features of clusters and droplets of SARS-CoV-2 N-protein with nucleic acid

LLPS is an entropically driven process, and whether the formation of a dense phase is promoted at higher or lower temperatures will reflect on the entropy gain or loss associated with the dense phase. The particle size will be a result of the free energies in both phases and the interfacial free energy (Berry et al., 2018; Vekilov, 2010), which likewise will have enthalpic and entropic contributions. Our ITC experiments of T_6_ binding N-protein revealed favorable entropic contributions from NA binding, which is thought to promote LLPS. To shed more light on this we carried out temperature scans to probe the energetics of LLPS of N-protein with different oligonucleotide ligands. For this we measured changes in particle size by DLS and microscopy, examined energetics in differential scanning calorimetry (DSC), in parallel to monitoring macromolecular solution structure reflected in CD and differential scanning fluorometry (DSF).

First looking at N-protein by itself, the magenta trace in **Figure 6A** shows the temperature dependent z-average (i.e., scattering intensity weighted) Stokes radius measured by DLS. A transition from the size of the N-protein dimer to a cluster of ≈60 nm occurs at 55°C, followed by a second transition to ≈100 nm particles at 65-70°C. Similar bimodal transitions are measured in differential scanning calorimetry (**Figure S5A**), with a strong endothermic peak already at a slightly lower temperature of 48°C, and a broad peak in the range of 60-70°C. DSF follows the same trends with a strong increase in the ratio of intrinsic fluorescence 350 nm to 330 nm at 50°C and a broader one at 65°C coinciding with the particle growth (**Figure 6B**). Notably, SARS-CoV-2 N-protein contains 5 Trp that are located exclusively in the NTD (3) or CTD (2), and likewise all 11 tyrosine residues are exclusively in these domains (7 in NTD and 4 in CTD), thus the structural changes reported by DSF can be attributed to the vicinity of the folded domains. By contrast, changes in the overall secondary structure are recorded in the CD thermal scans (**Figure 7ABC**). It shows a single transition at 46°C from the largely disordered state into a state with significantly reduced negative ellipticity at 200 nm and stronger ellipticity in the range of 220 - 230 nm, suggesting ≈10% increased helical content. To assess droplet formation during the CD temperature scan, far-UV absorbance traces were acquired simultaneously. At 95°C we measure a 2.2-fold increase at 200 nm with a spectrum characteristic of particle scattering (inset **Figure 7C**).

**Figure 6.**
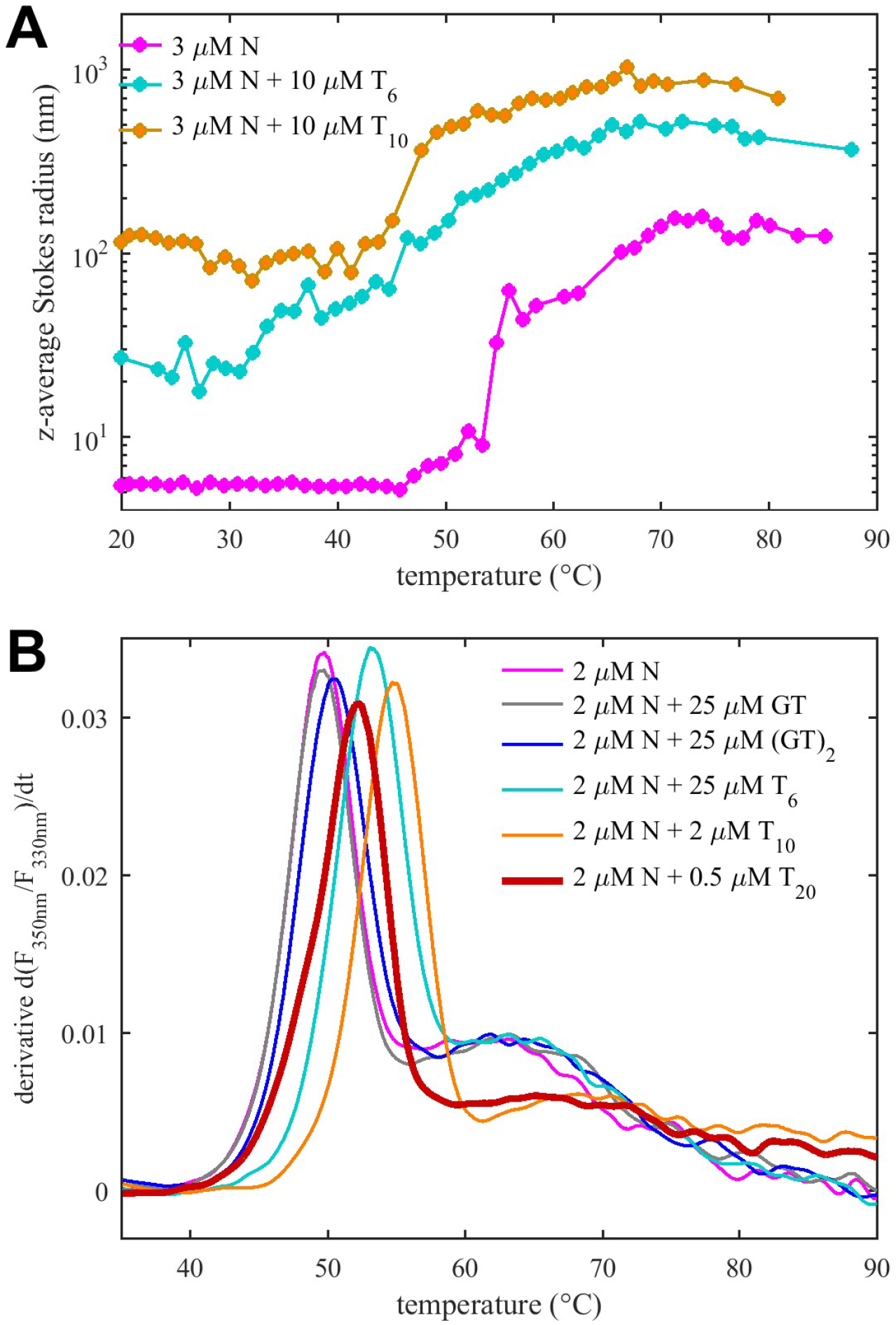
Temperature-Dependent Particle Size and LLPS. (A) Average Stokes radius of N-protein, with T_6_ and T_10_, as a function of temperature in DLS. (B) Temperature-derivative of the intrinsic fluorescence ratio at 350 nm to 330 nm (DSF) for N-protein with different ligands.

**Figure 7.**
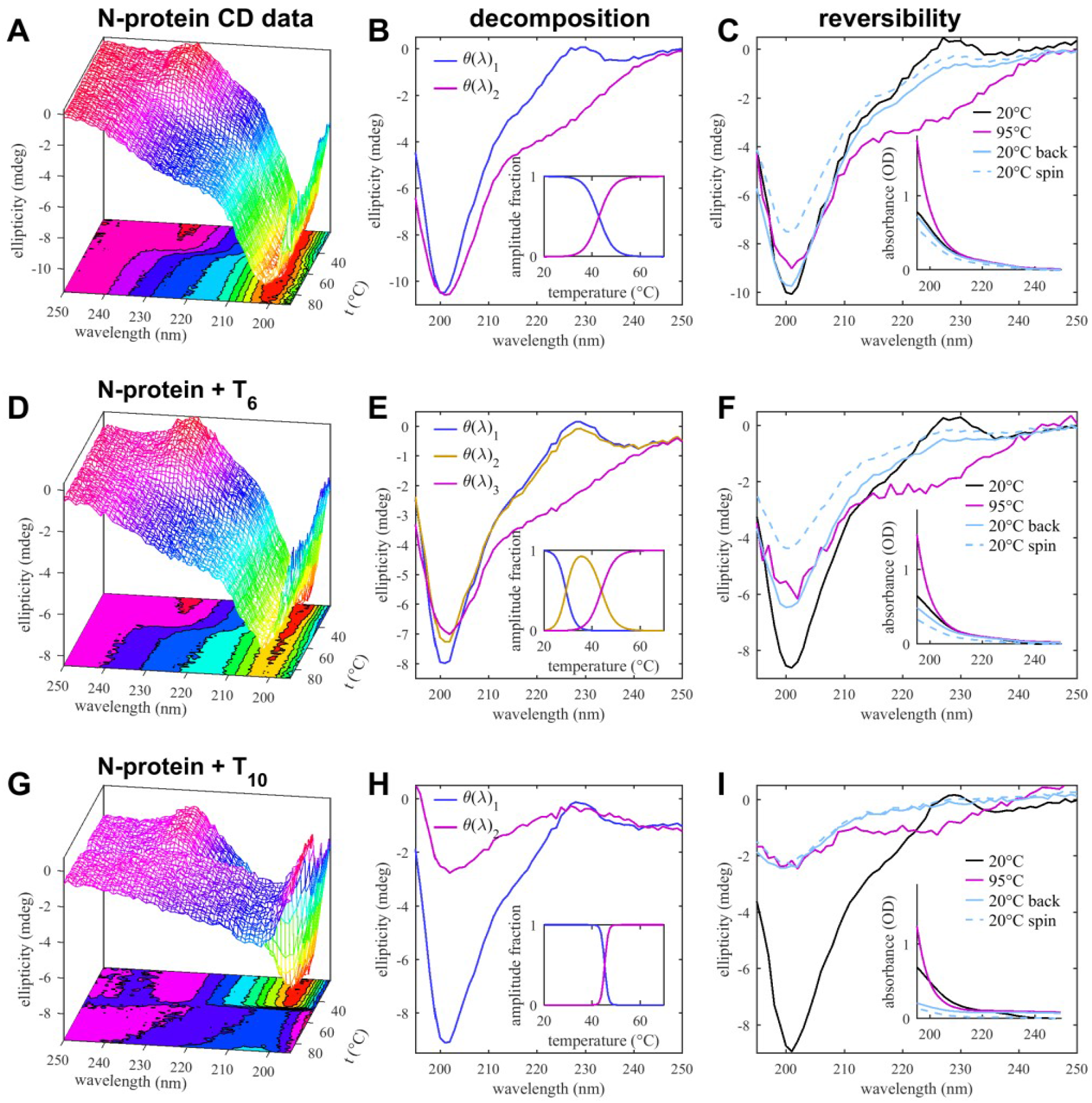
Temperature-Dependent Secondary Structure by CD. Top row (A-C) are data for 2 μM N-protein itself, middle row (D-F) for 2 μM N-protein with 10 μM T_6_, and bottom row (G-I) for 2 μM N-protein with 10 μM T_10_. The left column (A, D, G) shows raw data of CD spectra as a function of temperature. These data are decomposed into basis spectra shown in (B, E, H) and corresponding fractional amplitudes in the insets. The right column (C, F, I) shows reversibility by comparing initial spectra at 20°C, at 95°C, and those after returning the samples back to 20°C, before and after final spin to deplete undissolved condensates. The insets show corresponding reversibility of the far UV absorbance profiles as a measure of scattering contributions from clusters and droplets.

Remarkably, the structural transition observed in CD at higher temperature was largely reversible, as cooling the sample back to 20°C shows a return to the large negative ellipticity at 200 nm, and substantially reduced 220 nm ellipticity (cyan line in **Figure 7C**), along with a return of the absorbance spectrum close to the initial state. To quantify fractions of undissolved large particles at the end of the cooling process, we then applied a final 10 min spin in a tabletop centrifuge and reacquired the spectra. This led to a 20% reduction of both the CD and the absorbance signal amplitudes, indicating a minority fraction of undissolved large particles at the end of the cooling process.

Taken together, these data show that under our conditions of neutral pH and low salt, SARS-CoV-2 N protein can undergo a largely reversible endothermic transition to a more helically structured configuration that leads to the formation and accumulation of sub-microscopic particles or clusters. The latter exhibit a lag compared to the structural transition, which may be due to the difference in physical observables and growth kinetics.

In the presence of the short oligonucleotide T_6_, a similar picture arises with notable differences. By DLS (**Figure 6A**, cyan), already at 20°C the z-average Stokes radius is at ≈20-30 nm, i.e., significantly higher than the N-protein alone, which indicates the presence of traces of larger clusters resolved above (**Figure 5B**). There is continuous particle growth with an earlier midpoint at ≈50°C, producing particles of up to 500 nm radius. Even though the average Stokes radii appear to shrink above 70°C, this can be attributed to technical artifacts of particles outgrowing the size range of DLS. The CD scans of N-protein in the presence of excess T_6_ exhibit already at 20°C less negative 200 nm ellipticity than N-protein alone due to oligonucleotide binding, as described above. In the temperature scan (**Figure 7DEF**), this feature is exacerbated in a first transition at 31°C with an otherwise unchanged spectrum. This suggests enhanced T_6_ binding, consistent with the entropic contributions to binding seen by ITC above. At 47°C a second transition can be discerned, with the major change being the increased negative ellipticity in the range from 220-230 nm, resulting in a spectrum similar to that observed at higher temperature for N-protein alone. This spectral feature is again partially reversed upon cooling the sample, however, a larger fraction of signal (≈35%) is lost after a final spin, consistent with enhanced particle formation. Also, there is little indication of lag in the temperature scans between particle growth and structural transitions.

Qualitatively similar features, but even stronger particle growth and sharper transitions were observed for N-protein in the presence of the decanucleotide T_10_. The z-average particle size is ≈100 nm already at 20°C, and exhibits rapid growth at 48°C (**Figure 6A**). At the same temperature, the CD spectra reveal a single transition to strongly reduced 200 nm ellipticity with enhanced 220-230 nm ellipticity, similar to the high-temperature state seen with other oligonucleotide ligands, but in a much sharper transition mirrored also in DSC (**Figure S5B**). This suggests stronger cooperativity and/or faster kinetics of droplet formation. Spectral changes in CD are reversible, but with substantial signal loss after cooling down. Centrifuging the final sample makes little further difference in signal loss, suggesting that larger particles are formed that settle already under earth gravity. In fact, turbidity monitored in the far UV absorbance traces show a significant drop while heating up the cuvette, starting at the phase transition temperature (**Figure S6**), which demonstrates loss of material from the optical path.

As for the longest oligonucleotide T_20_, as described above, we observed the formation of very large particles already at room temperature, and these were therefore excluded from CD and DLS temperature scan experiments. However, due to the capillary format and more rapid temperature scans of DSF, mixtures of N-protein with T_20_ could be included in an experiment series comparing transitions as a function of oligomer length, suggesting similar major transitions ≈50-55°C (**Figure 6B**).

## Discussion

In the present work we report comprehensive biophysical characterization of the structure, energetics, and assembly states of N-protein and its complex states with NA oligomers of different lengths.

A basic question under current discussion is the solution state of purified full-length SARS-CoV-2 N-protein (Carlson et al., 2020; Cubuk et al., 2020; Ye et al., 2020). Our SV-AUC data unequivocally show N-protein in tightly bound and spatially extended dimers, under a wide range of ionic strength and temperature, consistent with SEC-MALS and SAXS data published by Zeng et al. (Zeng et al., 2020). Due to the extended shape these have larger hydrodynamic radii and would elute in size exclusion chromatography (SEC) at higher apparent molecular weights if compared to folded protein standards. Our findings appear to be in contrast to the observation of higher oligomers full-length SARS-CoV-2 N-protein in SEC-MALS reported by Ye at al. (Ye et al., 2020), but are consistent with NA-linked oligomerization reported in the same work (Ye et al., 2020). Linkage to NA binding may also explain higher oligomers reported for SARS-CoV-1 N-protein in experiments with unpurified N-protein from cell lysates (Cong et al., 2017; He et al., 2004b), as cellular RNA may be tightly bound even in fragments after nuclease treatment (Carlson et al., 2020). Subpopulations of higher-order oligomers have also been reported after chemical crosslinking, or in isolated domains (Chang et al., 2013; Luo et al., 2005; Tang et al., 2005; Yu et al., 2005; Zeng et al., 2020), which may point to ultra-weak transient higher oligomers evidenced also in our SV-AUC experiments. These may contribute to protein-protein interfaces amplified in NA-bound states (Chang et al., 2014).

Dimers are highly multi-valent for NA binding, as reported for SARS-CoV-1 (Chang et al., 2009). However, in the present study we found the binding mode and affinity depends strongly on the length of the oligonucleotide. The minimal length for measureable binding in our assays is 4 bases, with the capability to crosslink, or ‘scaffold’ N-protein arising at 20 bases.

Consistent with recent literature (Zeng et al., 2020; Zinzula et al., 2020), we find already a hexanucleotide is capable of high-affinity binding to N-protein, though only with 2 sites per dimer. Within the accessible concentration range in our work this produces well-defined complexes 2:2 without further assembly. This offers the opportunity to examine the effects of NA binding on N-protein structure and energetics in more detail. By calorimetry we find binding is both enthalpic and entropically driven. Since N-protein is highly basic at neutral pH, and NA binding is ionic strength dependent, the observed energetics is consistent with counter-ion release entropically contributing to binding (Ou and Muthukumar, 2006; Park et al., 2020). Simultaneously, we find that NA binding (hexanucleotides as well as longer oligonucleotides) is associated with reduced disorder of the N-protein chain as observed by CD spectroscopy. Furthermore, the NA-bound conformation is also more compact, as can be deduced from the smaller frictional ratio of the N-protein dimer once fully ligated with T_6_. Recently published cryo-EM data of SARS-CoV-2 N-protein in the presence of 7mers also showed 2:2 complexes (formed with similar affinities as measured here for T_6_), but suggested a rearrangement of the CTD dimer in cryo-EM relative to the crystal structure (Zinzula et al., 2020), highlighting N-protein flexibility and impact of NA-binding on conformation.

Decanucleotides bind with higher affinity than hexanucleotides, and this reveals at least four NA sites per dimer. When N-protein dimers are saturated with T_10_, dimer-dimer self-association can be observed, driven by higher protein concentrations. This is consistent with small dimer-dimer and tetramer-tetramer interfaces observed in crystal structures of SARS-CoV-1 N-protein that were hypothesized to be modulated by NA binding and part of the coassembly mechanism (Chang et al., 2014; Chen et al., 2007). Also, interactions at the extreme C-terminus (Ye et al., 2020) may come into play. Indeed, structural changes linked to protein-protein interactions have been speculated to exist based on the interpretation of LLPS experiments by Carlson and co-workers (Carlson et al., 2020).

With a 20nt NA ligand, we observe very high affinity binding, consistent with fluorescence polarization results by Wu et al. (Wu et al., 2020). Our data show that cross-linking of N-protein dimers by a single 20mer can occur. Spatially a 10mer is comparable in length to the NA-binding canyon of the NTD (Dinesh et al., 2020), and therefore the cross-linking ability of a 20mer but not the 10mer seems geometrically reasonable. This renders both N-protein and NA mutually multi-valent, and accordingly, we have observed even larger macromolecular complexes, consistent with an indefinite association process that leads to LLPS at significantly lower critical concentration and temperatures than shorter oligomers.

LLPS is an entropically driven demixing process that creates a liquid phase in droplets of high local macromolecular concentrations. While phase boundaries are commonly explored varying ligand concentrations, ionic strength, or pH (Cubuk et al., 2020; Perdikari et al., 2020; Savastano et al., 2020), we apply temperature changes as a driving parameter. This complements the recent study by Iserman et al. observing temperature-dependent LLPS by light microscopy (Iserman et al., 2020). Our biophysical data suggest that condensates form in two stages. In our experimental conditions of low-salt neutral pH buffer, N-protein alone undergoes LLPS only at elevated temperatures, and then produces only 60 – 100 nm particles. These clusters may represent nuclei for the growth of large droplets. Nuclei formation is enhanced by N-protein dimer interactions with NA; however, additional protein-protein and protein/NA interfaces are required for the formation of the larger droplets, consistent with the notion that multi-valency is an essential feature of LLPS (Reichheld et al., 2017). Multi-valency is further exacerbated by the multiple weak interaction sites of N-protein for NA, and accordingly we have measured 4-fold molar excess of T_10_ hosted in N-protein droplets. Our observation that binding of 20mer lowered the transition temperature (relative to LLPS with shorter NA) is in line with the observation by Iserman and co-workers that binding of longer genomic RNA lowers the transition temperature to 37°C (Iserman et al., 2020).

Interestingly, in contrast to other proteins undergoing LLPS (Reichheld et al., 2017), the phase separation of N-protein appears to be supported by structural transitions from disordered into a more ordered state, in configurational changes that are in addition to -- and distinct from -- those imposed by NA binding alone. This is reminiscent of the gain of structure reported by Zeng and co-workers from CD at 55°C (Zeng et al., 2020), and may arise from stabilization of transient helices identified in the disordered regions of SARS-CoV-2 N-protein (Cubuk et al., 2020). DSF results suggest that these changes also impact the folded domains (where the aromatic amino acids exclusively reside). In fact, our DSF traces of N-protein are very similar to those reported by Zinzula et al. for the isolated CTD (Zinzula et al., 2020). This may also reflect indirect contributions from changes in solvation in the proximity of the folded domains. In conclusion, while these experiments do not yet provide direct proof of the altered protein conformation once in the dense phase, we can speculate that this increased order persists, and that it promotes protein-protein interactions that, in conjunction with the higher collision frequency in the dense phase, support formation of well-defined ribonucleoprotein particles. The latter may proceed via preferred condensation of protein to select high-affinity recognition elements along the genomic RNA (Cubuk et al., 2020; Iserman et al., 2020), possibly in concert with M-protein interactions (Lu et al., 2020; Masters, 2019), and probably with architectural control of resulting structures being modulated by N-protein self-association properties (Joyeux, 2021).

The importance of LLPS *in vivo* in the viral life cycle is stressed by results from an evolutionary analysis of SARS-CoV-2 (Gussow et al., 2020), which found localization in nucleoli is correlated with increased pathogenicity and case fatality rate. LLPS may be regulated by small molecules, and thereby susceptible to small molecule inhibitors, some of which have already been identified in different laboratories (Carlson et al., 2020; Cubuk et al., 2020; Iserman et al., 2020; Jack et al., 2020; Zhao et al., 2020). We believe this strategy of therapeutic development will be supported by increasing detailed knowledge of molecular configurations and binding processes involved in N-protein functions.

## Acknowledgement

This work was supported by the Intramural Research Programs of the National Institute of Biomedical Imaging and Bioengineering, National Heart, Lung, and Blood Institute, National Cancer Institute, and National Institute of Neurological Disorders and Stroke, National Institutes of Health.

## Methods

### Protein and oligonucleotides

SARS-CoV-2 nucleocapsid protein (GenBank: BCA87368.1) expressed in *E. coli* with C-terminal His-tag, purified by Ni-chelating and size exclusion chromatography, was acquired from EXONBIO (catalog# 19CoV-N150, San Diego, CA). Compositional Mw is 46,981.06 Da, extinction coefficient at 280 nm is 43,890 M^−1^cm^−1^, and partial-specific volume if 0.717 ml/g. The protein was formulated in phosphate buffer pH 7.4, 250 mM NaCl and stored in frozen form. In our laboratory, we confirmed higher than 95% of purity from contaminating proteins by SDS-PAGE and SV-AUC (**Figure 1**). We verified the sequence by LC-MS/MS, and verified the protein mass by LC-MS, showing mono-dispersity 46851 ± 2 Da as expected after removal of initial methionine, in the absence of phosphorylation. The ratio of absorbance at 260 nm and 280 nm of the N protein stock was 0.516, confirming the absence of nucleic acids, as shown in the absorbance spectrum in **Figure S7**. (We initially surveyed six commercial sources, and found that many other suppliers had significant nucleic acid contamination, consistent with the observation by Iserman and co-workers (Iserman et al., 2020).) As described in the Results section, the N-protein exhibited secondary structures by CD and melting curves by DSF consistent with those reported in the literature. Depending on the purpose of the experiments, the N protein was dialyzed exhaustively against either PBS (Na_2_PO_4_ 10.1 mM, KH_2_PO_4_ 1.8 mM, KCl 2.7 mM, NaCl 137 mM, pH 7.4) or the low salt working buffer (Na_2_PO_4_ 10.1 mM, KH_2_PO_4_ 1.8 mM, KCl 2.7 mM, NaCl 10 mM, pH 7.4) prior to the subsequent biophysical characterization. In some experiments mentioned in the results the buffers were supplemented with surfactant P20 (catalog# BR100054, Cytiva, Marlborough, MA) at a final concentration of 0.005%. Final protein concentrations were measured by spectrophotometry, or in SV-AUC experiments by Rayleigh interferometry.

As described in the Results, some SV-AUC experiments were carried out with fluorescently labeled N-protein, which was produced with the Dylight488 amine-reactive antibody labeling kit (catalog# 53025, Thermo Fisher), using NHS ester chemistry by following the protocol provided by the vendor, achieving a labeling ratio of 0.498 based on UV-VIS spectrophotometry. Dylight488-labeled N protein was dialyzed against PBS. Fluorescent-detected SV-AUC experiments were carried out using bovine serum albumin (catalog# A7030, Sigma-Aldrich, St. Louis, MO) as a carrier in order to minimize surface adsorption of N protein in the AUC cells.

The oligonucleotides (GT) and (GT)_2_ were purchased from the Keck Biotechnology Resource Laboratory at Yale University (New Haven, CT), as purified with cartridge followed by lyophilization. T_6_ (TTTTTT), T_10_ (TTTTTTTTTT) and T_20_ (TTTTTTTTTT TTTTTTTTTT) were purchased from Integrated DNA Technologies (Skokie, IL), which were purified by HPLC and received in a lyophilized form. The oligonucleotides were reconstituted in PBS to reach a concentration of 500-1000 μM. Some of the oligonucleotides were subjected to dialysis against the working buffer (Na_2_PO_4_ 10.1 mM, KH_2_PO_4_ 1.8 mM, KCl 2.7 mM, NaCl 10 mM, pH 7.4) using the Spectrum™ Float-A-Lyzer™ G2 Dialysis Devices with MWCO 100-500 Da or 500-1000 Da, respectively. The final concentration was determined using a UV-Vis spectrophotometer and molar extinction calculated based on the sequences.

#### Analytical ultracentrifugation

SV experiments were carried out in a ProteomeLab XL-I analytical ultracentrifuge (Beckman Coulter, Indianapolis, IN) by following standard protocols (Schuck et al., 2015). Experiments with DyLight488-labeled N-protein were carried out in an Optima XL-A analytical ultracentrifuge equipped with a fluorescence detection system with excitation at 488 nm (AVIV Biomedical, Lakewood, NJ). Samples were loaded in 12- or 3-mm charcoal-filled Epon double-sector centerpieces with sapphire windows and loaded into an An-50TI 8-hole rotor. Unless otherwise mentioned temperature was equilibrated to 20°C, followed by acceleration to 50,000 rpm and data acquisition using Rayleigh interference optics and absorbance optics at 230 nm, 260 nm, and/or 280 nm. Data analysis was carried out using the software SEDFIT (National Institutes of Health) and to calculate the sedimentation coefficient distribution c(s) (Schuck, 2016). Integrated weight-average sedimentation coefficients were assembled into isotherms and fitted with binding models in the software SEDPHAT (National Institutes of Health) (Schuck and Zhao, 2017).

#### Dynamic light scattering

Autocorrelation data were collected in a NanoStar instrument (Wyatt Technology, Santa Barbara, CA). 100 μL samples at 3 μL N-protein in the presence or absence of oligonucleotides were inserted into a 1 μL quartz cuvette (WNQC01-00, Wyatt Instruments), using excess sample to prevent evaporation in the observation chamber. Laser light scattering was measured at 658 nm at a detection angle of 90°. For the temperature scans, a ramp rate of 1°C/min was applied with 5 sec data acquisitions and averaging 3 replicates for each temperature point. Data were collected and processed by using software Dynamics 7.4 (Wyatt Instruments) or SEDFIT (National Institutes of Health).

#### Isothermal titration microcalorimetry

ITC titrations were performed using an ITC200 microcalorimeter (Malvern, Northampton, MA) at 20°C. Protein and oligonucleotides samples were dialyzed against working buffer, diluted to the target concentrations, and degassed. 48 μM N-protein was loaded into the cell, and 200 μM T_6_ was loaded into the syringe, and after equilibration 17 injections of 2.25 μL were made in intervals of 300 sec. The power trace was integrated using NITPIC (Keller et al., 2012) and analyzed using ITC models of SEDPHAT in global multi-method modeling (Zhao and Schuck, 2012; Zhao et al., 2015).

#### Circular dichroism spectroscopy

CD spectra were acquired in a Chirascan Q100 (Applied Photophysics, U.K.). Samples were measured in 1 mm pathlength cells, with 1 nm steps and 1 sec integration time. Results are averages of 3 acquisitions. Backgrounds of corresponding buffers (with or without oligonucleotides, respectively) were subtracted. For temperature scans, data were acquired in 1 nm intervals with integration times of 0.5 sec, without repeats, applying a temperature ramp rate of 1°C/min. The Global3 software from Applied Photophysics was used to deconvolute the multi-wavelengths temperature scans. Helicity fractions were estimated using the software BeStSel (Micsonai et al., 2018).

#### Widefield microscopy

Images were acquired on a Nikon EclipseTE2000 U microscope equipped with a PCO Edge 4.2 LT camera (PCO AG, Kelheim, Germany) with sensor dimensions of 2048×2048. Images were collected using a 100X 1.4 numerical aperture oil objective lens and a bin factor of 4 to produce 512×512 images with a pixel size of 156 nanometers as determined with a calibrated graticule. The transmitted light source was a collimated white light LED (Thorlabs, Inc., Newton, NJ) passed through a green interference filter. 20 μL samples were added to a glass cover slip for imaging at room temperature. For reaction mixtures, N protein and T_20_ were combined and mixed immediately prior. Imaging was started immediately and continued for 5 min with 5 frames per sec. To observe the effect of dilution, the same amount of working buffer was added to the mixture on the coverslip and gently mixed by pipetting, followed by imaging.

#### Differential scanning fluorometry

DSF was carried out using a Tycho instrument (Nanotemper, Germany). 10 μL samples were loaded in capillaries (TY-C001, Nanotemper). The intrinsic protein fluorescence was measured at 350 nm and 330 nm, and the first derivative of the intensity ratio was calculated as a function of temperature. The temperature ramp rate was 30°C/min and data were acquired from 35 to 95 °C.

#### Differential scanning calorimetry

DSC was performed in a VP-DSC instrument (Malvern Panalytical, U.K.) using standard protocols. 500 μL of 10 μM N-protein samples with or without oligonucleotide and corresponding buffers were loaded in the sample and reference cell, respectively. A temperature ramp rate of 1°C/min was used.

