## Supplemental Figures for "Energetic and structural features of SARS-CoV-2 N-protein co-assemblies with nucleic acids"

**Figure S1: N-Protein Dimer Dissociation**

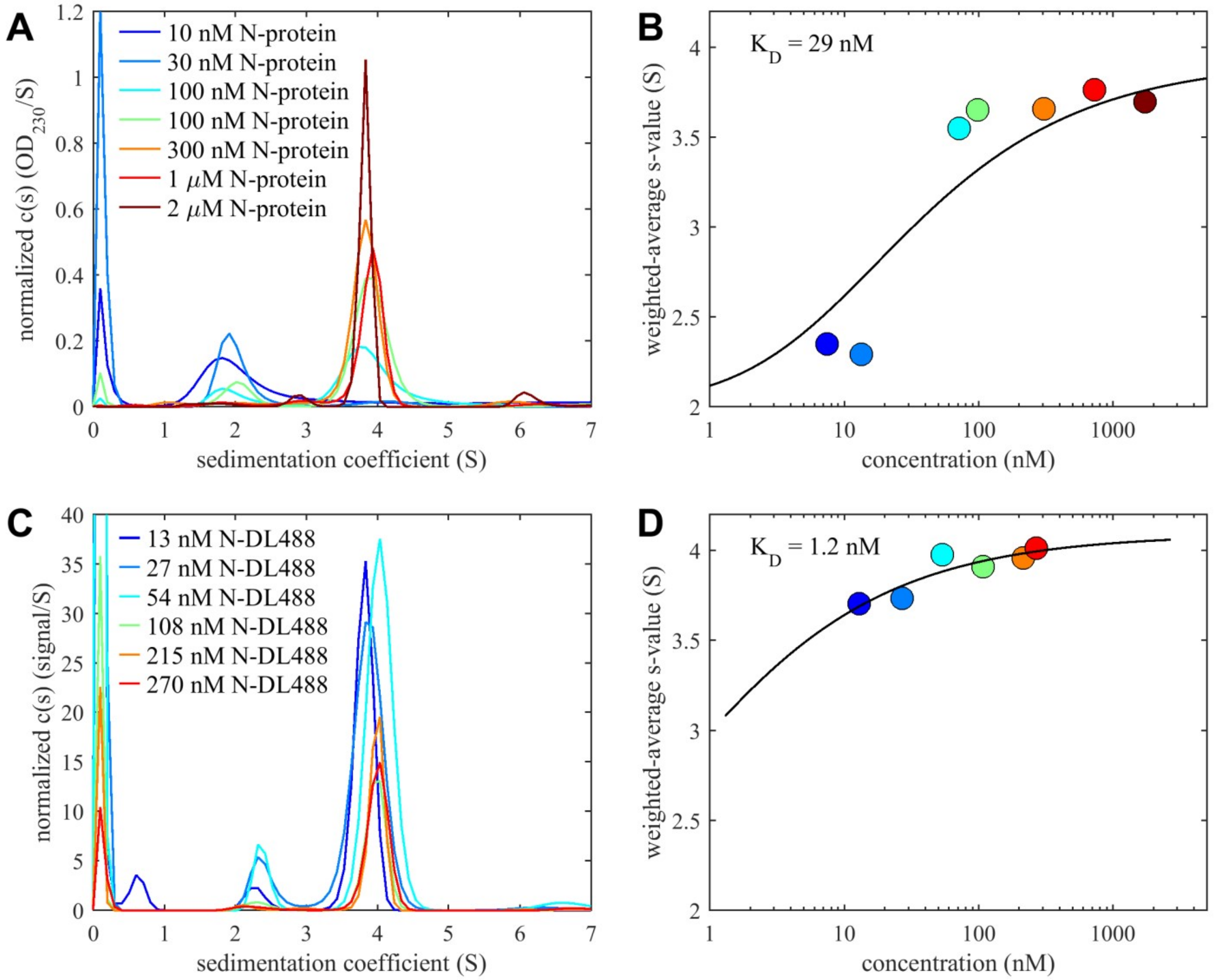


(A) Sedimentation coefficient distributions of N-protein in PBS supplemented with 0.005% surfactant P20, obtained from SV-AUC experiments using far-UV detection at 230 nm. Shown are traces labeled with nominal loading concentrations (including a replicate at 100 nM).

(B) Isotherm of weight-average s-values of N-protein with P20 shown in Panel A, and best-fit monomer-dimer model. Concentrations are determined from integrated sedimentation boundary amplitudes.

(C) Sedimentation coefficient distributions of DyLight488-labeled N-protein in PBS based on SV-AUC using fluorescence detection (MacGregor et al., 2004; Zhao et al., 2013).

(D) Isotherm of weight-average s-values of DyLight-488-labeled N-protein in Panel C, and best-fit monomer-dimer model.

**Figure S2: Circular Dichroism Spectra of N-protein with surfactant P20 or with fluorescent tag**

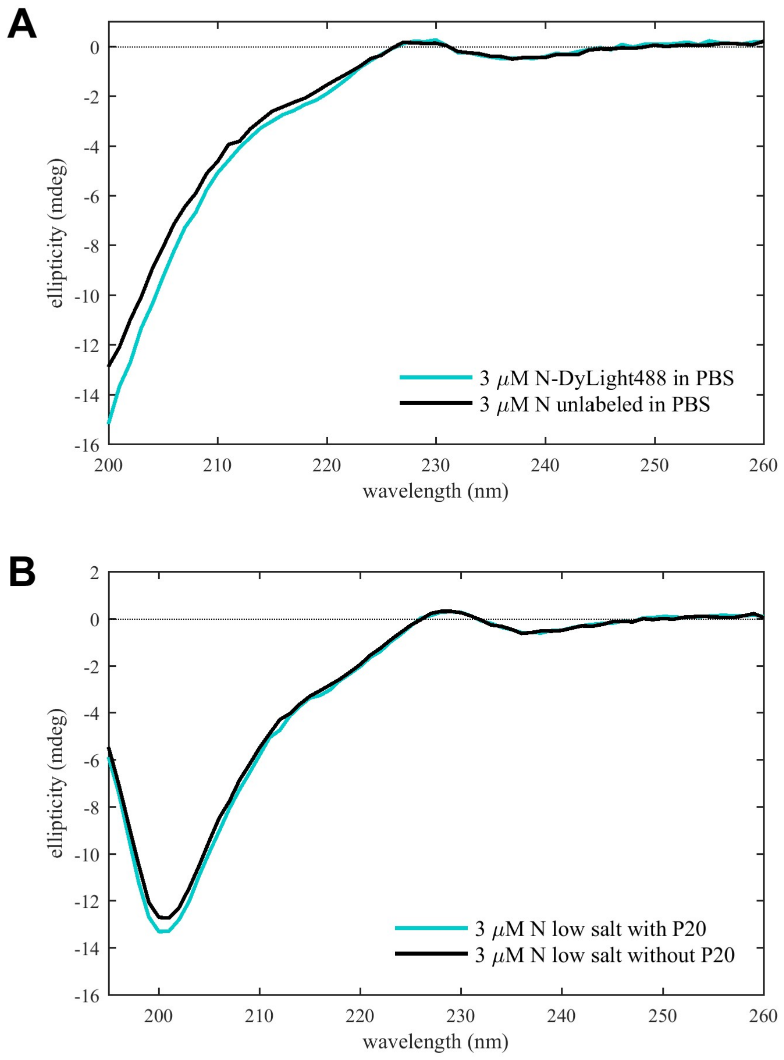


(A) Comparison of CD spectra in PBS of unlabeled and DyLight488-labeled protein.

(B) Comparison of CD spectra of unlabeled N-protein in 12 mM $KH_2PO_4/Na_2HPO_4$, 3 mM KCl, 10 mM NaCl, pH 7.4 and in the same buffer supplemented with 0.005% P20.

**Figure S3: Oligonucleotide binding to N-protein in high ionic strength buffer**

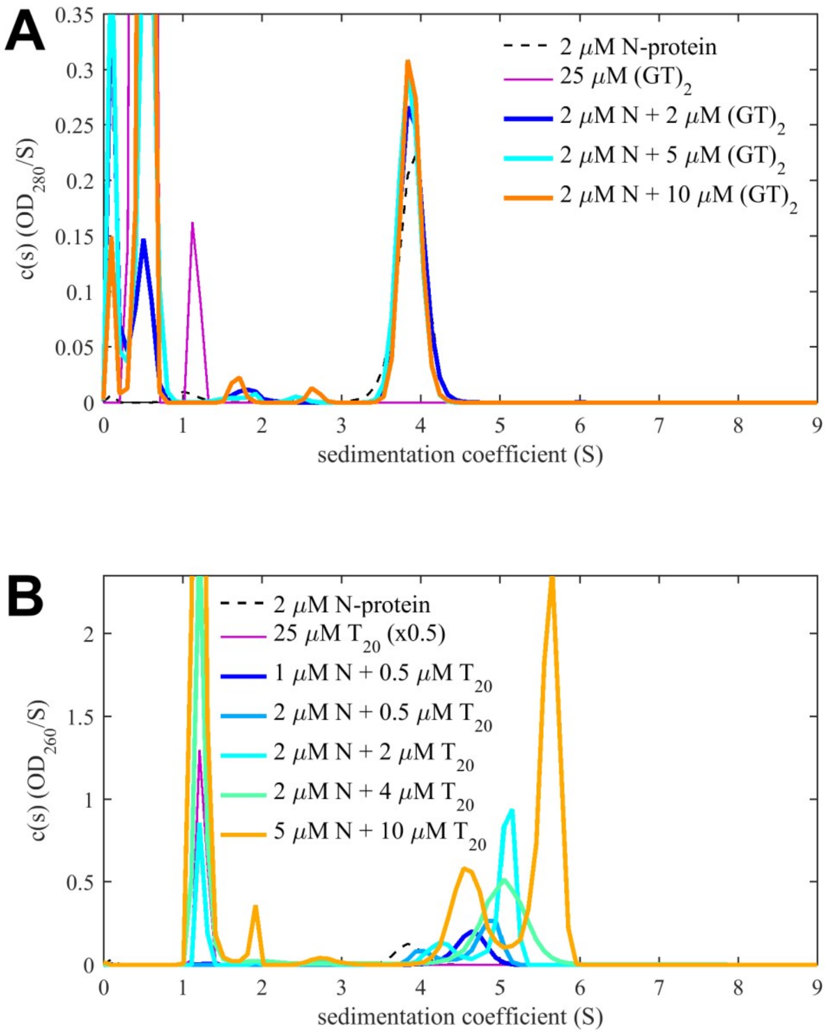


(A) Sedimentation coefficient distributions of mixtures of N-protein and $(GT)_2$ in PBS show no increase in s-value and no significant increase in boundary amplitudes with $(GT)_2$ concentration.

(B) Concentration series of N-protein with $T_{20}$ in PBS shows significant binding, but at strongly reduced level than in low salt buffer, exhibiting lower complex sedimentation coefficients.

**Figure S4: Reversibility of droplet formation of N-protein with $T_{20}$.**

Widefield microscopy image of 5 µM N-protein with 10 µM $T_{20}$ before (A) and after (B) twofold dilution.

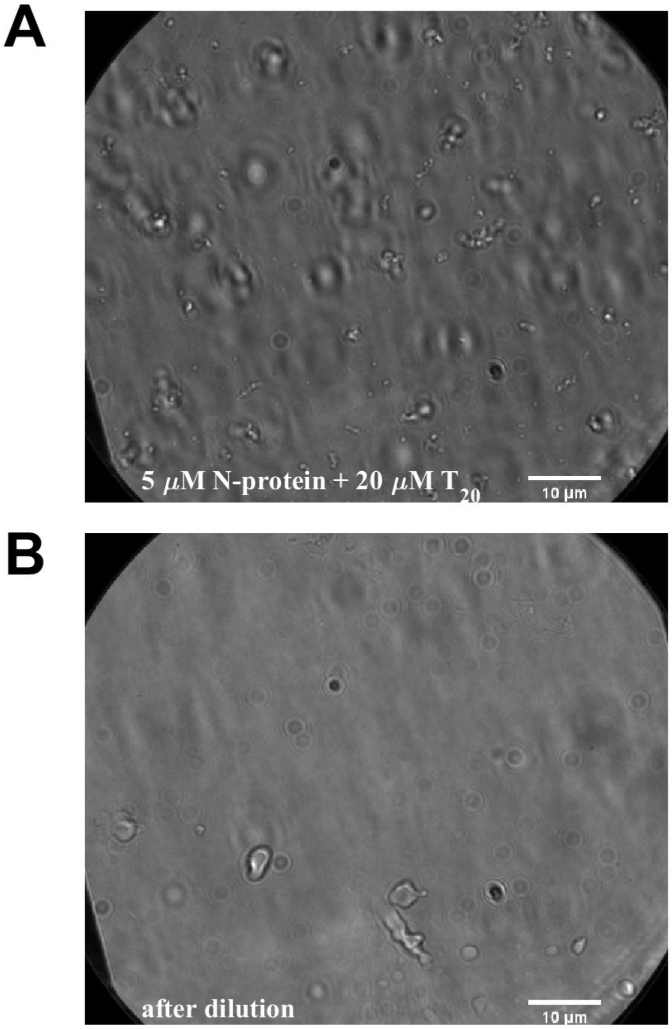

**Figure S5: Differential scanning calorimetry of N-protein with $T_{10}$.**

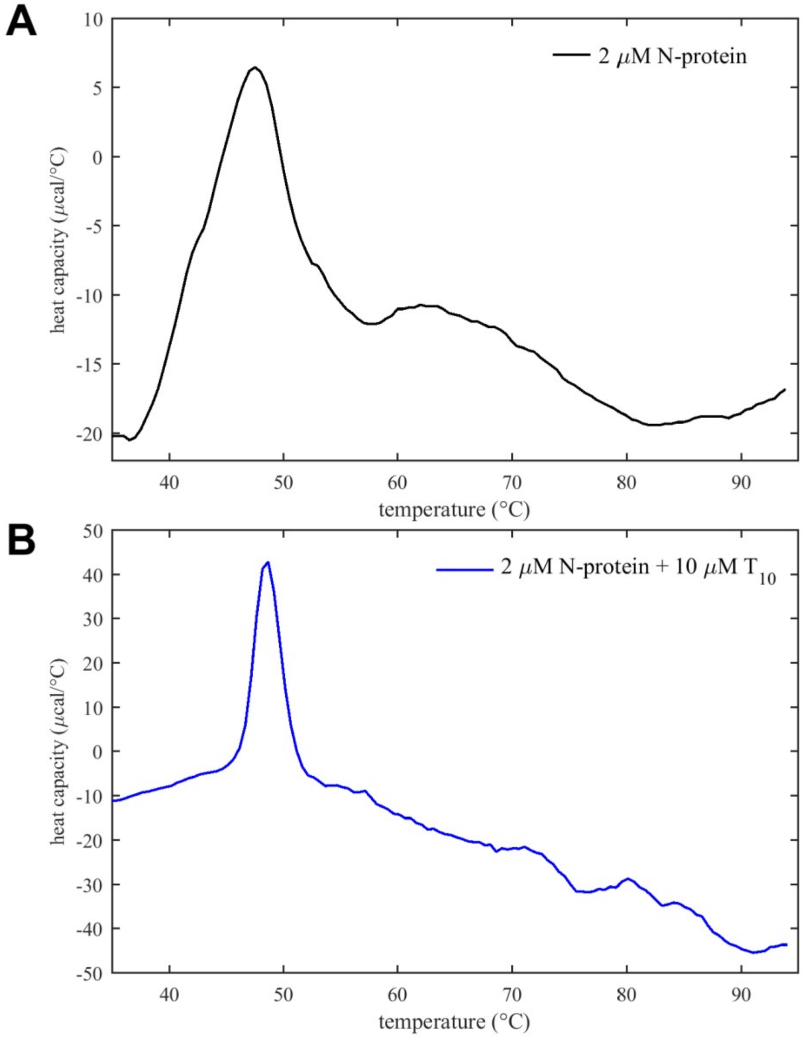

**Figure S6: Far-UV absorbance during temperature scan**

Absorbance traces at 200 nm recorded during CD temperature scans in **Figure 7**.

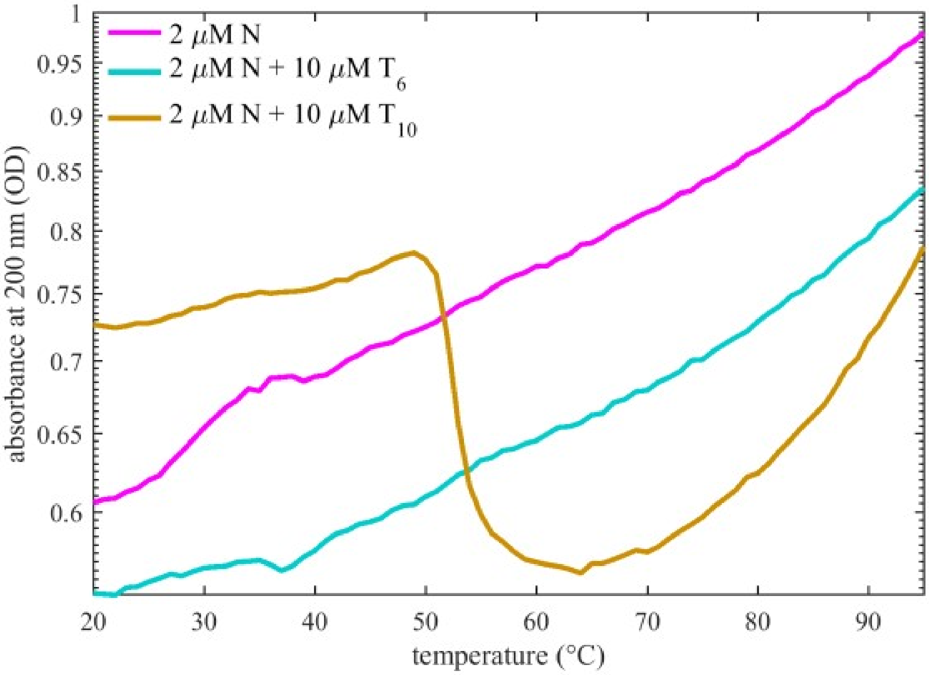

**Figure S7: Absorbance spectrum of N-protein**

Shown is a 1:10 dilution of N-protein stock solution after dialysis against working buffer.

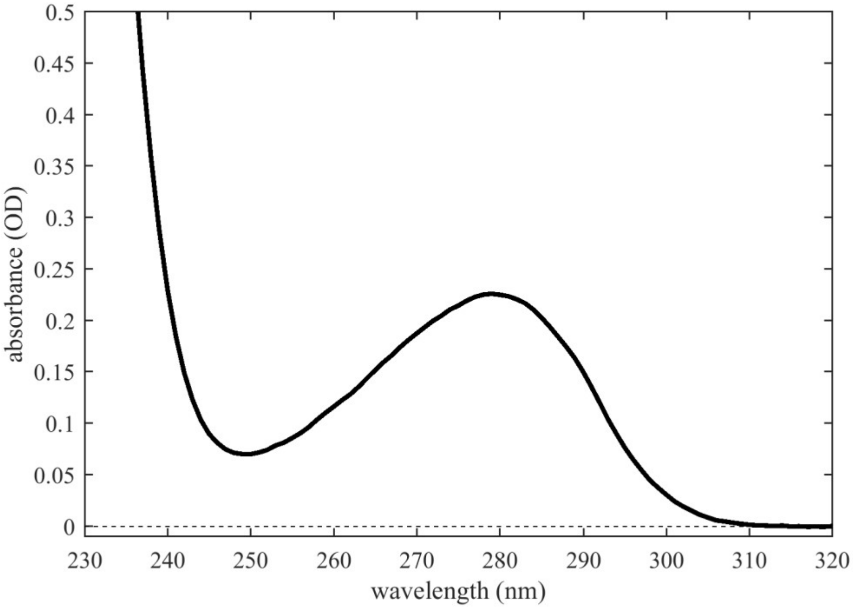
